# Conserved roles of RECQ-like helicases Sgs1 and BLM in preventing R-loop associated genome instability

**DOI:** 10.1101/119677

**Authors:** Carolina A. Novoa, Emily Yun-Chia Chang, Maria J. Aristizabal, Yan Coulombe, Romulo Segovia, Yaoqing Shen, Christelle Keong, Steven J.M. Jones, Jean-Yves Masson, Michael S. Kobor, Peter C. Stirling

## Abstract

Sgs1 is a yeast DNA helicase functioning in DNA replication and repair, and is the orthologue of the human Bloom’s syndrome helicase BLM. Here we analyze the mutation signature associated with *SGS1* deletion in yeast, and find frequent copy number changes flanked by regions of repetitive sequence and high R-loop forming potential. We show that loss of *SGS1* increases R-loop accumulation and sensitizes cells to replication-transcription collisions. Accordingly, in *sgs1*Δ cells the genome-wide distribution of R-loops shifts to known sites of Sgs1 action, replication pausing regions, and to long genes. Depletion of the orthologous BLM helicase from human cancer cells also increases R-loop levels, and R-loop-associated genome instability. In support of a direct effect, BLM is found physically proximal to DNA:RNA hybrids in human cells, and can efficiently unwind R-loops *in vitro*. Together our data describe a conserved role for Sgs1/BLM in R-loop suppression and support an increasingly broad view of DNA repair and replication fork stabilizing proteins as modulators of R-loop mediated genome instability.

## INTRODUCTION

Genome instability is an enabling characteristic of tumor formation because it creates mutational diversity in pre-malignant cell populations and thereby allows the necessary mutations in driver genes to occur at a sufficiently high frequency (Stratton et al., 2009). One of the best understood ways in which genome instability arises in cancer is an acquired defect in a DNA repair pathway. Mutations in homologous recombination (HR), nucleotide excision repair (NER), crosslink repair, and mismatch repair (MMR) are all clearly linked to an increased risk of cancer (Curtin, 2012). Germline mutations in DNA repair dramatically increase cancer risk and can be associated with other symptoms, while somatically acquired cancer driver mutations in the same repair pathways are found in sporadic cancers.

One such DNA repair proteins is the RECQ-like helicase BLM which is involved in resolution of concatenated DNA molecules during HR, at replication forks, and in anaphase (Bohm and Bernstein, 2014). BLM can act on a wide array of substrates *in vitro* including Holliday junctions, D-loops, G-quadruplexes, DNA:RNA hybrids, and single-stranded overhangs (Croteau et al., 2014; Popuri et al., 2008). Germline BLM mutations lead to Bloom’s syndrome, which is characterized by cancer predisposition, short stature and other symptoms (de Renty and Ellis, 2017). At the cellular level defects in BLM are characterized by high levels of sister chromatid exchanges, and DNA replication and mitotic defects (Bohm and Bernstein, 2014). BLM is highly conserved across evolution, and much of what is known about its function was first described for its orthologue in *Saccharomyces cerevisiae*, Sgs1. Research on Sgs1 has linked its activity to homologous recombination, both at the stage of end-resection and double-Holliday junction resolution, restart of stalled DNA replication forks, meiosis and telomere maintenance (Ashton and Hickson, 2010). Both BLM and Sgs1 have multiple interacting partners that regulate their activity and cooperate with them in catalyzing DNA transactions. BLM/Sgs1 forms a complex with Topoisomerase III and RMI1/2, which work together with BLM to decatenate DNA molecules. The BLM-Top3-Rmi1/2 complex further associates with members of the Fanconi Anemia pathway to help process stalled DNA replication forks for example during interstrand crosslink repair (Ling et al., 2016; Suhasini and Brosh, 2012).

Recently, defects in DNA repair proteins have been linked to a novel mechanism of genome instability involving the formation of excessive DNA:RNA hybrids on genomic DNA. These hybrids form a structure called an R-loop, in which RNA binds to a complementary DNA strand and exposes the non-template strand as a ssDNA loop (Chan et al., 2014b). R-loops are thought to cause genome instability primarily by interfering with DNA replication. R-loop collision with replication forks leads to fork stalling and an increase in double-strand breaks or error prone mechanisms of replication (Chan et al., 2014b). The best understood players in R-loop metabolism are those involved in RNA processing where normal transcript elongation, termination, splicing, packaging, nuclear export and RNA degradation have all been shown to suppress R-loop formation (Gomez-Gonzalez et al., 2011; Li and Manley, 2005; Mischo et al., 2011; Stirling et al., 2012; Wahba et al., 2011). Interestingly, defects in canonical DNA repair proteins such as HR factors BRCA1 and BRCA2 (Bhatia et al., 2014; Hatchi et al., 2015; Wahba et al., 2013), NER proteins XPG and XPF (Sollier et al., 2014), the Fanconi Anemia pathway (Garcia-Rubio et al., 2015; Schwab et al., 2015) and the DNA damage response kinase ATM (Tresini et al., 2015) have all been associated with stabilization or signaling involving R-loops. Moreover, R-loops have been shown to contribute to DNA replication stress in these DNA repair mutants and in some cases a direct role for the repair protein in R-loop removal has been suggested (e.g. R-loop displacement by the FANCM helicase (Schwab et al., 2015)). Indeed, the BLM protein is known to cooperate with the HR pathway and is critical for the activation of the Fanconi Anemia pathway. Moreover, Sgs1 has synthetic phenotypes with RNaseH2 deletions in yeast, suggesting a functional cooperation between the two proteins (Chon et al., 2013; Kim and Jinks-Robertson, 2011).

In this study we began by assessing the accumulated mutation spectrum associated with *SGS1* deletion in diploid yeast. This analysis revealed a signature of copy-number alterations associated with homeologous flanking sequences. These breakpoint regions were correlated with regions of very high R-loop occupancy and, which coupled with recent literature led us to investigate the role of Sgs1 and BLM in R-loop suppression. We show that in yeast deleted for *SGS1* genome instability is partially R-loop dependant, and that R-loops accumulate at sites of Sgs1 action in the genome. We go on to confirm R-loop associated genome instability in BLM depleted human cells, showing that BLM localizes near R-loops in cells and is capable of resolving R-loops as efficiently as D-loops *in vitro*. Together these data establish Sgs1/BLM as a regulator of R-loop coupled genome instability, adding to the growing repertoire of DNA repair proteins with functions in R-loop mitigation, and extending the notion that transcription-replication collisions are one of the drivers of mutagenesis in cancer.

## RESULTS

### Genome instability in sgs1Δ yeast occurs at transcribed repetitive regions

In order to determine the genomic locations of instability in *sgs1*Δ cells we performed a mutation accumulation (MA) and whole-genome sequencing (WGS) experiment (Stirling et al., 2014). Passaging homozygous *sgs1*Δ/Δ diploids for approximately 1000 generations created a set of 12 MA strains that we sequenced at >50x coverage (**Fig. 1A**). As a control, we conducted the same experiment with WT and cells lacking *MUS81*, a structure-specific endonuclease which also works in the processing of replication forks and HR intermediates (Boddy et al., 2001; Hanada et al., 2007). This analysis revealed a modest ^∼^2-fold increase in single-nucleotide variants (SNVs) for *sgs1*Δ/Δ and a smaller increase for *mus81*Δ/Δ as compared to WT, which were similar to the rates seen at the *CAN1* reporter locus (Segovia et al., 2017).

**Figure 1.**
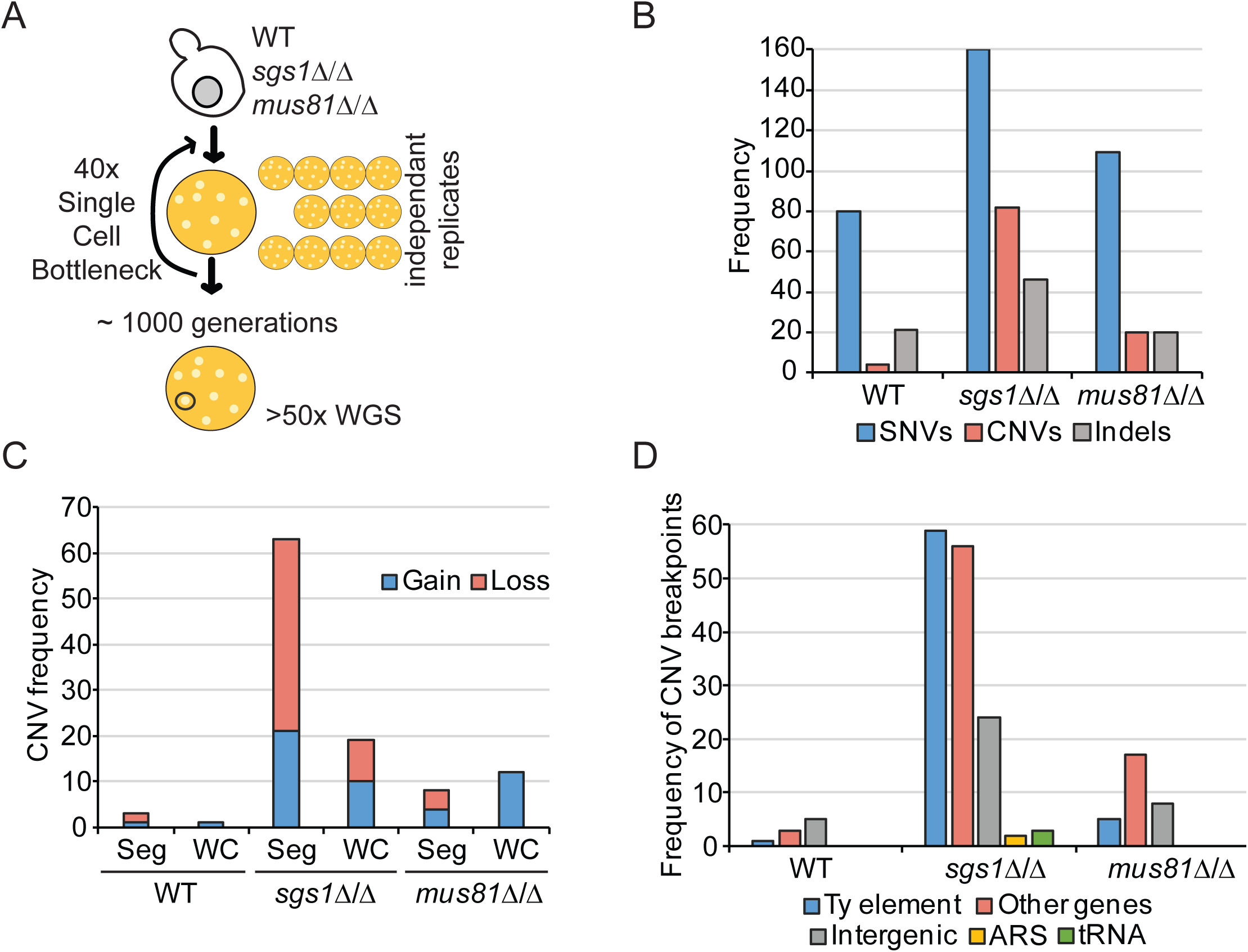
The mutation spectrum of *SGS1* deficient yeast. (A) Schematic of the 1000 generation MAWGS approach. (B) Frequency of SNVs, CNVs and indels in MA lines of the indicated strains. (C) CNV breakpoint classes in MA lines. WC = whole chromosome aneuploidy; Seg = segmental copy number change. (D) CNV breakpoint characteristics in MA lines. For more detailed information see **Table S1**.

However, both strains had a large increase in predicted copy-number variants (CNVs), including a ^∼^12x increase for *sgs1*Δ/Δ compared to WT (**Fig. 1B**). In addition, *sgs1*Δ/Δ cells displayed more segmental (Seg) CNVs, whereas *mus81*Δ/Δ showed proportionally higher whole chromosome (WC) aneuploidy (**Fig. 1C**). Analysis of the predicted breakpoints shows that most originate within or near a TY retrotransposon element (**Fig. 1D**; **Table S1**). Indeed, Sgs1 has been ascribed a role in Ty1 element expansion based on its role in HR (Bryk et al., 2001). Other breakpoints appear to be at telomeres, and at a set of protein coding genes. Breakpoint-associated genes were often pairs of paralogous genes (e.g. *ENA1/ENA2; PHO3/PHO5; FCY21/FCY22; ALD2/ALD3*, etc. **Table S1**) in the yeast genome and this, along with the TY element enrichment, is consistent with the role for Sgs1 in rejecting homeologous recombination reactions that could lead to CNVs between repetitive or paralogous sequences (Myung et al., 2001). Ty1 elements are among the most highly transcribed features in the yeast genome, and have previously been implicated as hotspots of R-loop formation in yeast by DNA:RNA immunoprecipitation (DRIP) (Chan et al., 2014a; El Hage et al., 2014; Wahba et al., 2016). Telomeric CNVs, which likely reflect terminal deletions, are consistent with a known role for Sgs1 in promoting telomere replication (Hardy et al., 2014). Although, telomeres are also sites of high R-loop formation due to TERRA transcription (Balk et al., 2013).

To explore the correlation between sites of instability in *sgs1*Δ and R-loops we assessed R-loop occupancy from previously published DRIP profiles at breakpoint-associated protein coding genes. Remarkably, R-loops signal was significantly higher than the average background of published WT DRIP peaks (10.2 (n =19) versus 4.1 (n=14307); Mann-Whitney Test p < 0.0001) (Chan et al., 2014a). Overall, the mutation signature of *sgs1*Δ/Δ cells supports its known specific role in promoting non-crossover events and rejecting homeologous recombination (Myung et al., 2001). Given both the recent linkage of fork protection factors such as BRCA2 and other Fanconi Anemia proteins to R-loops (Bhatia et al., 2014; Garcia-Rubio et al., 2015; Schwab et al., 2015), and that regions of instability are associated with reported hotspots of R-loop formation we decided to investigate a potential link of Sgs1/BLM to R-loops.

### Loss of SGS1 exhibits synergistic genome instability with R-loop suppressors

Yeast Sgs1 has been ascribed a downstream or cooperative role with RNaseH2 in suppressing genome instability, presumed to be related to shared roles in DNA replication and potential cooperation at sites of transcription mediated instability (Chon et al., 2013; Kim and Jinks-Robertson, 2011). To assess the fitness consequences of a putative R-loop increase in *sgs1*Δ cells we first determined if combining *SGS1* deletion with mutation in additional R-loop regulators in the THO complex (*MFT1*), RNaseH enzymes (*RNH1* and *RNH201*) and senataxin (*SEN1*) resulted in synergistic fitness defects (Gomez-Gonzalez et al., 2009; Huertas and Aguilera, 2003; Mischo et al., 2011; Skourti-Stathaki et al., 2011). Loss of Sgs1 significantly exacerbated fitness defects in *rnh201*Δ, *sen1-1* and *mft1*Δ cells (**Fig. 2A**). These defects were enhanced in *sen1-1* and *rnh201*Δ in the presence of hydroxyurea (HU) which causes DNA replication stress (**Fig. S1**). We next examined genome instability using the A-like Faker (ALF) assay for disruption of the MAT locus in chromosome III (Stirling et al., 2011). Double mutants of *SGS1* and the THO complex subunits *MFT1* or *THP2* exhibited dramatic increases in the frequency of chromosome III loss. Similarly, mutations in *SEN1* or deletion of *RNH1/201* lead to large increases in ALF frequency (**Fig. 2B**). Analysis of a plasmid based *LEU2* direct repeat recombination (Prado et al., 1997) also revealed synergistic increases in recombination frequency when combining deletion of *SGS1* and either *MFT1* or *THP2* (**Fig. 2C**). These synergies are not surprising since loss of Sgs1 is known to promote hyperrecombination through loss of its role in resolution of Holliday junctions (Ashton and Hickson, 2010). R-loop frequency increases with transcript frequency and length in direct repeat recombination assays. Therefore, to implicate Sgs1 further in transcription-associated recombination we assessed the roles of transcript length and frequency using derivatives of the *LEU2* direct repeat systems and comparing to *mft1*Δ for length and transcription-dependent instability. While *sgs1*Δ had higher recombination frequencies in all assays, comparing the rates of recombination in a short transcript (L) and a long transcript (LYΔNS) plasmid system revealed only a 1.5 fold increase in recombination in WT, but a 6-fold increase in *sgs1*Δ (**Fig. 2D**). Similarly, shifting the galactose inducible GL-LacZ recombination cassette (Gonzalez-Aguilera et al., 2008) from dextrose to galactose led to 678x increase in recombination in WT, but a 1277x increase in *sgs1*Δ (**Fig. 2E**). These data extend and support previous observations of transcription-associated instability in *SGS1* mutants and functional synergy between Sgs1 and Rnh201 (Chon et al., 2013; Kim and Jinks-Robertson, 2011).

**Figure 2.**
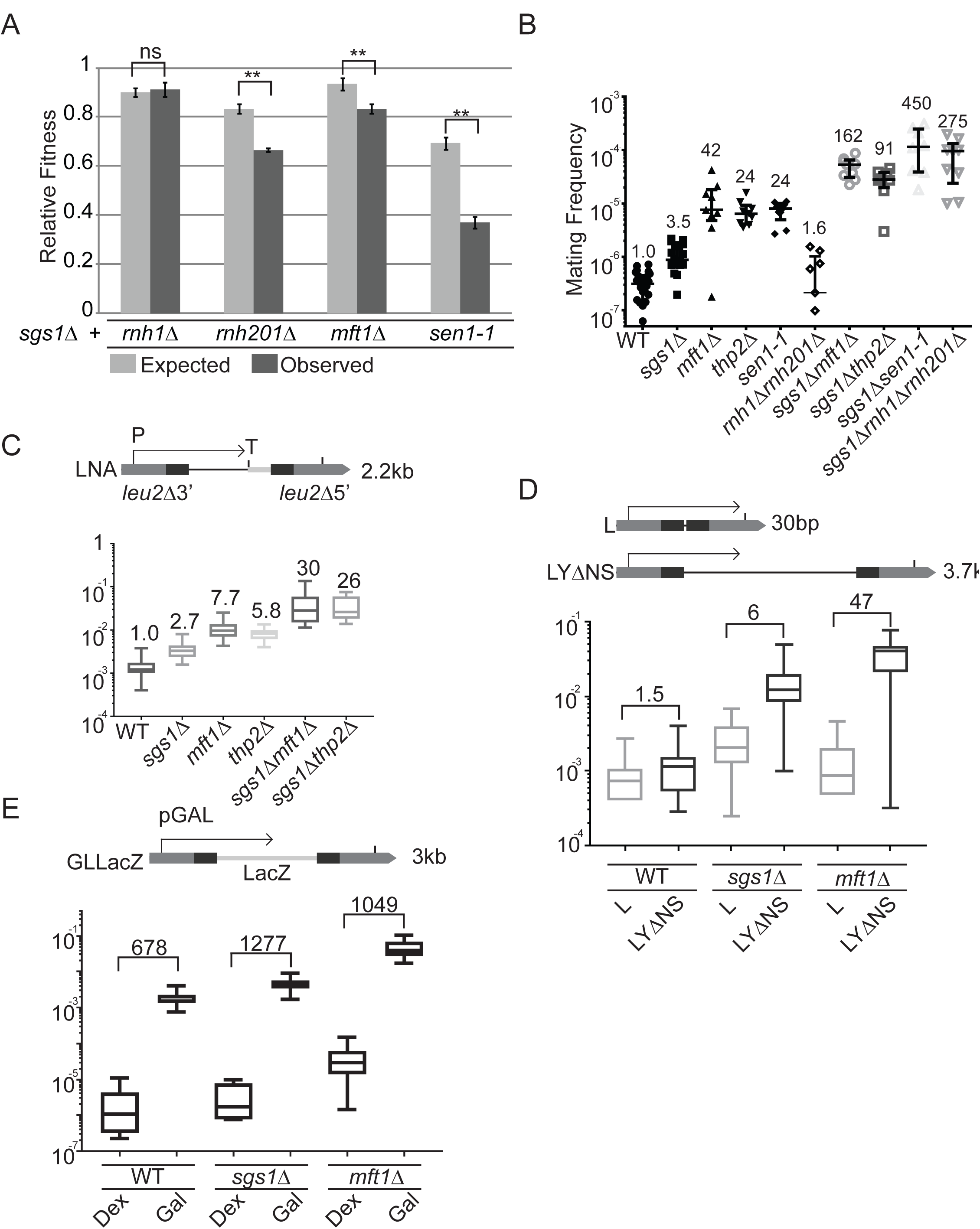
Cooperative genome maintenance by R-loop regulators and Sgs1. (A) Growth defects in strains lacking Sgs1 and the indicated gene. Quantitative growth curve analysis showed observed fitness values were significantly lower than expected values. (B) Mating frequency as a measure of chromosome instability by the a-like faker assay (Duffy et al., 2016). Fold increase over WT is shown above each measurement in the indicated strains. (C-E) Plasmid based *LEU2* recombination frequency measurements in the indicated strains. Above each panel is a schematic of the assay which measure recombination frequency, transcript length dependence and transcript frequency dependence, respectively. The length of the intervening sequence between the *leu2* repeats (dark bars) is shown at right. The fold increase over WT is shown above each bar. The direction of transcription is indicated by an arrow. For E, Dex = dextrose (low expression) and Gal = galactose (high expression). For all figures exact p-values are listed or asterisks indicate: * = p<0.05, ** = p<0.01, *** = p<0.001, ****= p<0.0001.

### R-loop formation and consequences in sgs1Δ

To directly assess the levels of DNA:RNA hybrids in *sgs1*Δ genomes, we used chromosome spreads and quantified staining with the S9.6 monoclonal antibody. This analysis showed a reproducible increase in S9.6 staining compared to WT (**Fig. 3A**). R-loops drive transcription-replication conflicts leading to recombination. To determine the effect of Sgs1 on this process we tested recombination rates in a plasmid system where transcription is driven only in S-phase by a promoter that is oriented to be colliding or co-directional with the origin of replication (*i.e*. IN or OUT respectively; **Fig. 3B**). This analysis showed that *sgs1*Δ driven hyper-recombination was significantly enhanced when S-phase (HHF) transcription is colliding with DNA replication but was unaffected by controls where transcription occurs in G1 (CLB-IN) or G2 (BLB-IN) phase (**Fig. 3B**). We next determined whether spontaneous DNA damage in cells lacking *SGS1* could be R-loop dependent. While deletion of *SGS1* leads to a small increase in levels of DNA damage in the genome as measured by Rad52-YFP foci, preventing reliable analysis of suppression, combined loss of RNaseH2A (i.e. *sgs1*Δ*rnh201*Δ) leads to a strong increase in DNA damage (**Fig. 3C**) (Chon et al., 2013). Importantly, this effect was suppressed by overexpression of RNaseH1. Since RNaseH1 only degrades RNA in DNA:RNA hybrids, as opposed to functioning in ribonucleotide excision repair, this shows that the synergistic DNA damage in *sgs1*Δ*rnh201*Δ cells is due to R-loops. It is important to acknowledge that cells lacking *SGS1* exhibit hyper-recombination in scenarios independent of R-loops and based on its well-established role in homologous recombination and replication fork protection, only a subset of genome instability events must be related to R-loop stabilization.

**Figure 3.**
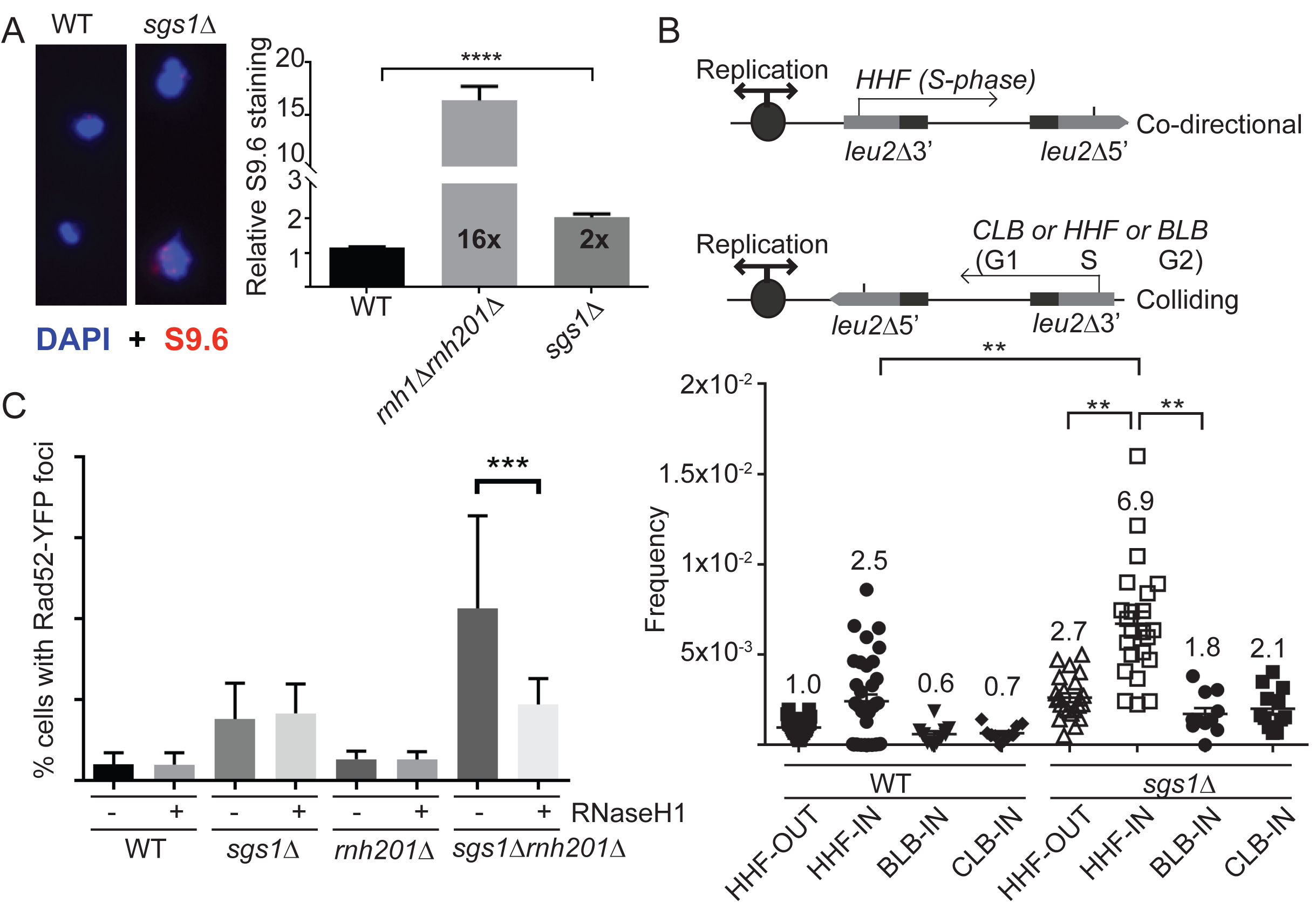
Transcription-replication conflicts and R-loops in *sgs1*Δ cells. (A) S9.6 staining for DNA:RNA hybrids in yeast chromosome spreads. Left, representative images; Right quantification of signal intensity per nucleus. Fold increase over WT is indicated in each bar. (B) Hyper-recombination due to replication-transcription collisions. Schematics of transcription direction and cell cycle stage of each promoter are indicated above. Below quantification of recombination frequencies for the indicated strain and plasmid. Fold increases over WT +*HHF-out* are shown above each bar. (C) DNA damage synergy in *sgs1*Δ*rnh201*Δ is R-loop dependant. Quantification of Rad52-YFP foci in the indicated strain with either an empty vector (-) or a Gal-inducible RNH1 (+) construct. The increase in foci in *sgs1*Δ*rnh201*Δ suppressed by overexpression of *RNH1*.

### SGS1 deletion shifts the profile of R-loops genome-wide

Given the increase in DNA:RNA hybrid levels in chromosome spreads and observed relationship with transcription-associated recombination, we hypothesized that loss of *SGS1* would alter the landscape of R-loops across the yeast genome. To test this we profiled the positions of DNA:RNA hybrids using an established S9.6 immunoprecipitation and microarray protocol (DRIP-chip) (Chan et al., 2014a). *SGS1* deletion lead to an overall increases in R-loop occupancy with a particular effect at sites of known Sgs1 action. Specifically, rDNA, TY retrotransposons, and telomeres, exhibited significantly increased R-loop levels in *sgs1*Δ compared to WT (**Fig. 4A-4C** and **Fig. S2** and **Fig. S3**). At rDNA, where Sgs1 is known to play a role in promoting both replication and transcription (Mundbjerg et al., 2015), we observed significantly enhanced instability in *sgs1*Δ which could be suppressed by ectopic expression of RNaseH1 (**Fig. 4D**). rDNA instability appeared to decrease in RNaseH1-expressing WT cells, consistent with the reported role for R-loops at this locus (El Hage et al., 2010), but did not reach significance.

**Figure 4.**
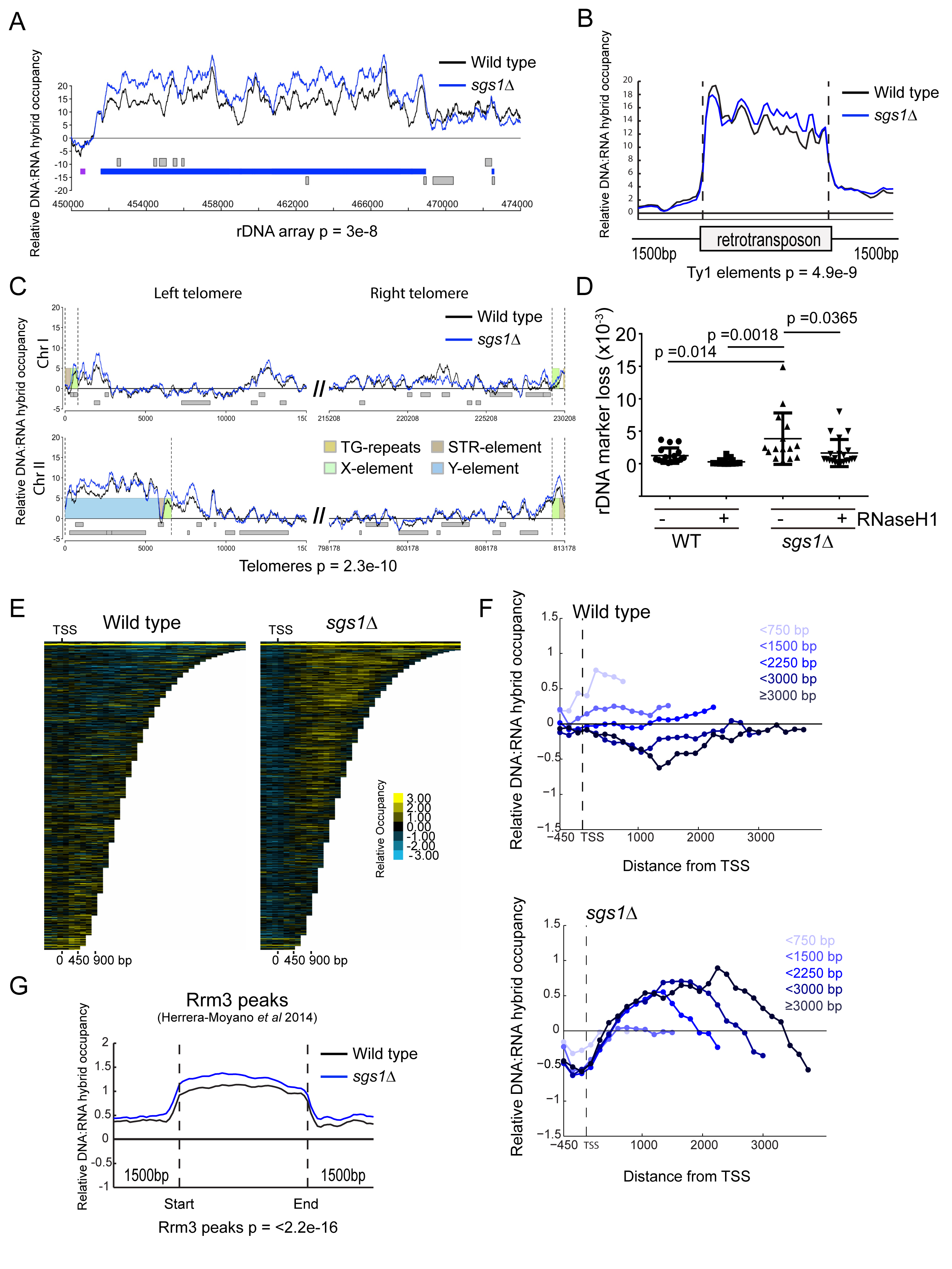
R-loops accumulate at sites of Sgs1 action in the yeast genome. (A) DNA:RNA occupancy at the rDNA locus. The MAT score is shown over the chromosomal coordinates of the region in bp (x-axis). (B) Averaged DNA:RNA occupancy over all Ty1 retrotransposons in the yeast genome. (C) Representative telomere occupancy of DNA:RNA hybrids. For A-C the p-value associated with increased occupancy in *sgs1*Δ versus WT is shown below the figure. (D) rDNA instability assay. Rate of loss of the *URA3* gene from the rDNA is indicated. P-values associated with significant differences are indicated and all other comparisons were not significant (ANOVA with Tukey’s test). (E) Chromatra plots showing a heat map of DNA:RNA occupancy over protein coding genes sorted by length and aligned at the TSS (transcription start site)(Hentrich et al., 2012). (F) Average genome-wide DNA:RNA occupancy in WT and *sgs1*Δ as a function of gene length. WT shows decreasing occupancy in longer gene length bins, while *sgs1*Δ shows the opposite trend. Mean values and statistics are reported in the main text. (G) Average DNA:RNA hybrid occupancy across previously identified Rrm3 peaks (Herrera-Moyano et al., 2014) for the indicated strains. p-value comparing increased occupancy in *sgs1*Δ is noted below.

Analysis of R-loop occupancy at mRNA encoding genes using CHROMATRA (Hentrich et al., 2012) illustrates an overall shift toward R-loop signal at long genes in *sgs1*Δ cells (**Fig. 4E**). Analysis of the 205 genes (**Table S2**) significantly occupied by DRIP-chip signal in both replicates of *sgs1*Δ but not WT, confirmed a significant shift toward longer than average genes (i.e. *sgs1*Δ DRIP genes were 1706bp, compared to 1338bp for all genes; Mann-Whitney p=0.0156). We analyzed the occupancy of DNA:RNA hybrids at all genes in gene size bins (**Fig. 4F**), confirming a trend toward longer genes in *sgs1*Δ compared to WT. The distribution of genes with increased DRIP signal in *sgs1*Δ also showed a bias to subtelomeric regions (**Fig. S2**), as 20% of *sgs1*Δ-specific DRIP genes (41 of 205) fell within 10kb of a telomere compared with just 2.7% of all annotated genes (Fisher test, p<0.0001) (**Fig. S2**). Moreover, if we remove subtelomeric genes within 10kb of the chromosome end, the average length of the remaining 164 *sgs1*Δ-specific DRIP increases significantly to 1967bp versus the genome-average of genes >10kb from ends (Mann-Whitney p<0.0001). This supports that gene length is independently driving increased DRIP in *sgs1*Δ. These observations are consistent with our finding that transcript length increases recombination frequency in *sgs1*Δ cells (**Fig. 3**). In addition, we reanalyzed genes associated with CNV breakpoints in our mutation accumulation experiment (**Fig. 1**) and find that breakpoint genes are also significantly longer than the genome average (1745bp compared with 1338bp, Mann-Whitney p=0.0425).

Finally, we tested the correlation between DRIP signal and previously reported occupancy of the Rrm3 helicase (Santos-Pereira et al., 2013), which has been associated with difficult to replicate regions and is required for robust growth in *sgs1*Δ cells (Schmidt and Kolodner, 2004). This analysis revealed a correlation between DRIP and Rrm3 occupancy in WT cells that was significantly enhanced in the *sgs1*Δ DRIP profiles (**Fig. 4G**). Together, this supports a model in which Sgs1 reduces R-loops at Rrm3 binding sites and thereby normally helps to mitigate transcription-replication conflicts and genome instability.

### R-loops accumulate and cause DNA damage in BLM depleted cell lines

The human orthologue of Sgs1 is the Bloom’s syndrome helicase BLM. To determine whether a role of Sgs1 in R-loop metabolism is conserved in mammalian cells, we used siRNA to target BLM in HeLa cells (**Fig. 5A**), and measured R-loop levels by immunofluorescence. Knockdown of BLM leads to a significant increase in S9.6 staining, which could be abolished by transfecting GFP-RNaseH1into the cells (**Fig. 5B-C**). Similar staining results were obtained for BLM-/-knockout derivatives of the near diploid HCT116 cell line compared to an isogenic control line (**Fig. S4**). Like Sgs1 in yeast, BLM depletion increases genome instability phenotypes and we therefore examined the effects of RNaseH1 overexpression in BLM knockdown cells on genome instability. BLM knockdown caused chromosome instability as measured by increased micronucleus formation after 48 hours (**Fig. 5D**), likely as a result of an inability to resolve anaphase bridges at mitosis (Naim and Rosselli, 2009). This increase in micronuclei formation was significantly suppressed by overexpression of GFP-RNaseH1 (**Fig. 5D** and **5E**). BLM knockdown also induced DNA breaks as measured by the neutral comet assay (**Fig. 5F** and **5G**) and ectopic expression of GFP-RNaseH1 significantly reduced these BLM-knockdown-induced increases in comet tail moment. Together these data suggest that a significant proportion of DNA damage and genome instability in BLM-depleted mammalian cells may be due to R-loops.

**Figure 5.**
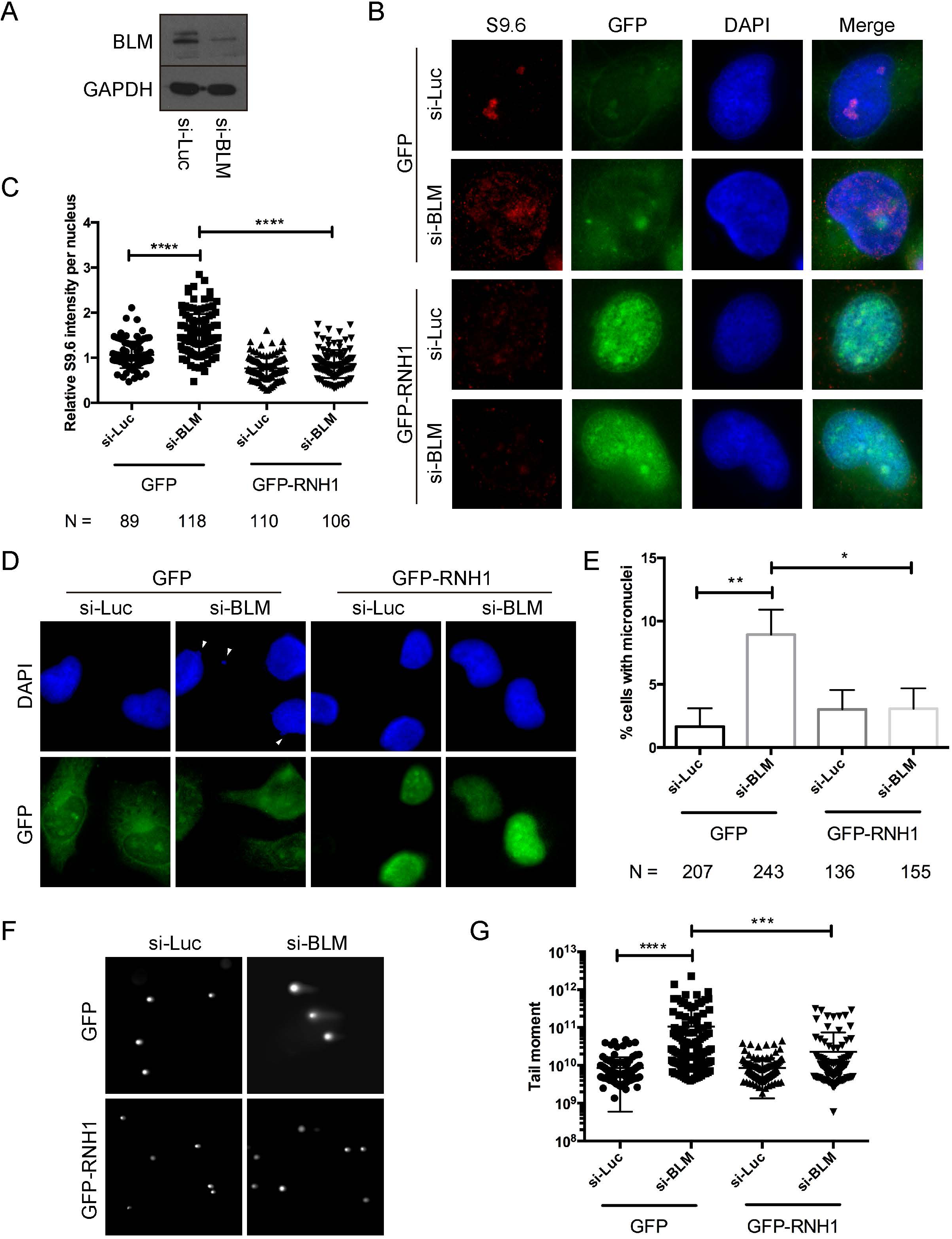
R-loop accumulation and DNA damage in BLM-depleted cells. (A) Effective knockdown of BLM in HeLa cells. Western blot showing BLM levels compared to a GAPDH control after siRNA targeting BLM or a control (Luc). (B) Representative images of S9.6 staining in cells treated with the indicated siRNA and transfected with either a control vector (GFP) or one expressing GFP-RNaseH1. (C) Quantification of S9.6 signal intensity for nuclear area in the indicated conditions. Cells scored across 3 independent replicates are noted below. (D) RNaseH-dependent chromosome instability in BLM deficient HeLa cells. Representative images of micronuclei (white arrowheads) in si-BLM treated cells expressing GFP or GFP-RNaseH1. (E) Quantification of micronuclei frequency in the indicated conditions. Cells counted are indicated below. (F) RNaseH-dependent DNA breaks in BLM deficient HeLa cells. Representative comet tail images from single-cell electrophoresis. (G) Quantification of comet tail moment under the indicated conditions. T-tests were used for comparisons shown.

### Assessing direct effects of BLM on R-loops

There are several possible mechanisms by which Sgs1 and BLM could impact R-loop mediated genome instability, including effects on replisome stability at transcription-replication conflicts, or through direct unwinding of R-loops, alone or collaboratively with the Fanconi Anemia (FA) pathway. To support these direct models, and rule out potential indirect effects, for example, through effects on DNA damage signaling pathways or transcriptional effects (Grierson et al., 2012; Tresini et al., 2015), we tested BLM helicase activity directly. While BLM has previously been shown to unwind R-loops *in vitro* (Popuri et al., 2008), we measured this activity in comparison to an efficiently unwound D-loop substrate and found that purified BLM unwound R-loops and D-loops of the same sequence with nearly identical efficiencies and that this occurred in an ATP and concentration dependent manner (**Fig. 6A**). To determine whether this could be occurring in cells, we performed a proximity ligation assay (PLA) with antibodies targeting BLM and DNA:RNA hybrids. Remarkably, BLM showed a clear and reproducible PLA signal in cells with S9.6 that was significantly higher than in single primary antibody controls, thus showing that BLM comes in close proximity to DNA:RNA hybrids in cells (**Fig. 6B**). Finally, to interrogate potential synergy with the FA pathway, we measured hybrid accumulation in single or double knockdown cells. As expected knockdown of FANCD2 induced R-loops (Garcia-Rubio et al., 2015; Schwab et al., 2015), but this increase was epistatic to coincident knockdown of BLM (**Fig. 6C**). Furthermore, BLM was required for HU induced increases in FANCD2 foci (**Fig. S5**), supporting literature placing these factors in the same pathway during replication stress (Ling et al., 2016; Panneerselvam et al., 2016). These data support to a model where local effects of BLM at R-loops occur in the same pathway as those of FANCD2. Together, our data support a conserved mechanism for Sgs1/BLM in suppressing transcription-associated genome instability at R-loop sites (Fig. 7).

**Figure 6.**
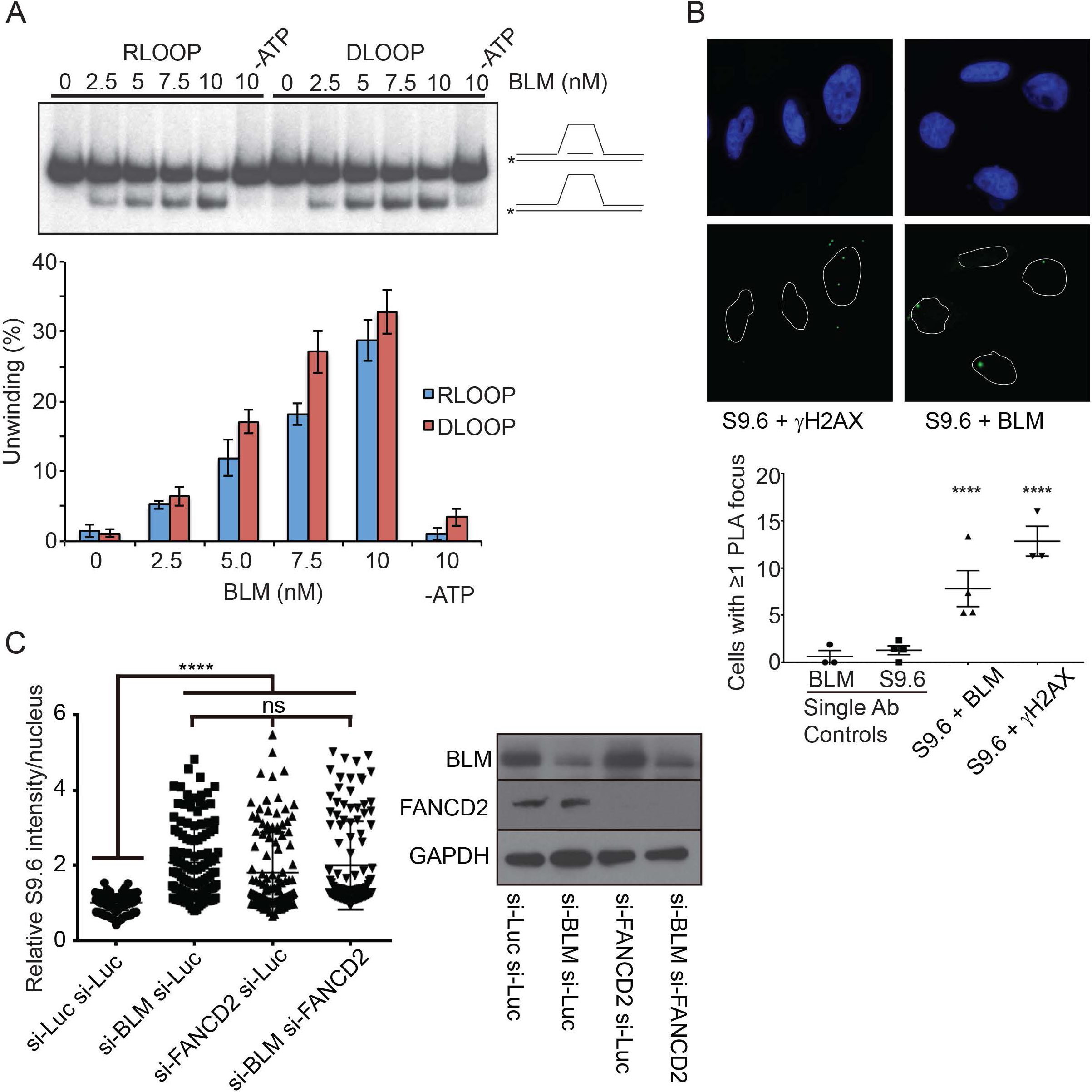
Potential mechanism of BLM-dependant R-loop mitigation. (A) Comparative R-loop and D-loop unwinding by BLM. Schematics of the slow migrating oligonucleotide loop substrate and faster migrating product are shown at right. BLM concentration is listed above. Quantification of unwinding efficiency is shown below. (B) Proximity ligation of BLM and DNA:RNA hybrids in cells. Representative images of PLA signal in HeLa cells in the presence of the indicated primary antibodies. Quantification of cells with PLA signal across replicates are shown below. S9.6 and γH2AX were previously associated in cells using PLA and serve as a positive control(Stork et al., 2016). Pooled count data for single primary antibody controls and dual antibody PLA reactions were compared using a Fisher exact test. (C) Nuclear S9.6 staining data for the indicated treatments. Quantification is shown at left, and western blot to confirm double knockdown efficiency is shown at right. ns = not significant.

**Figure 7.**
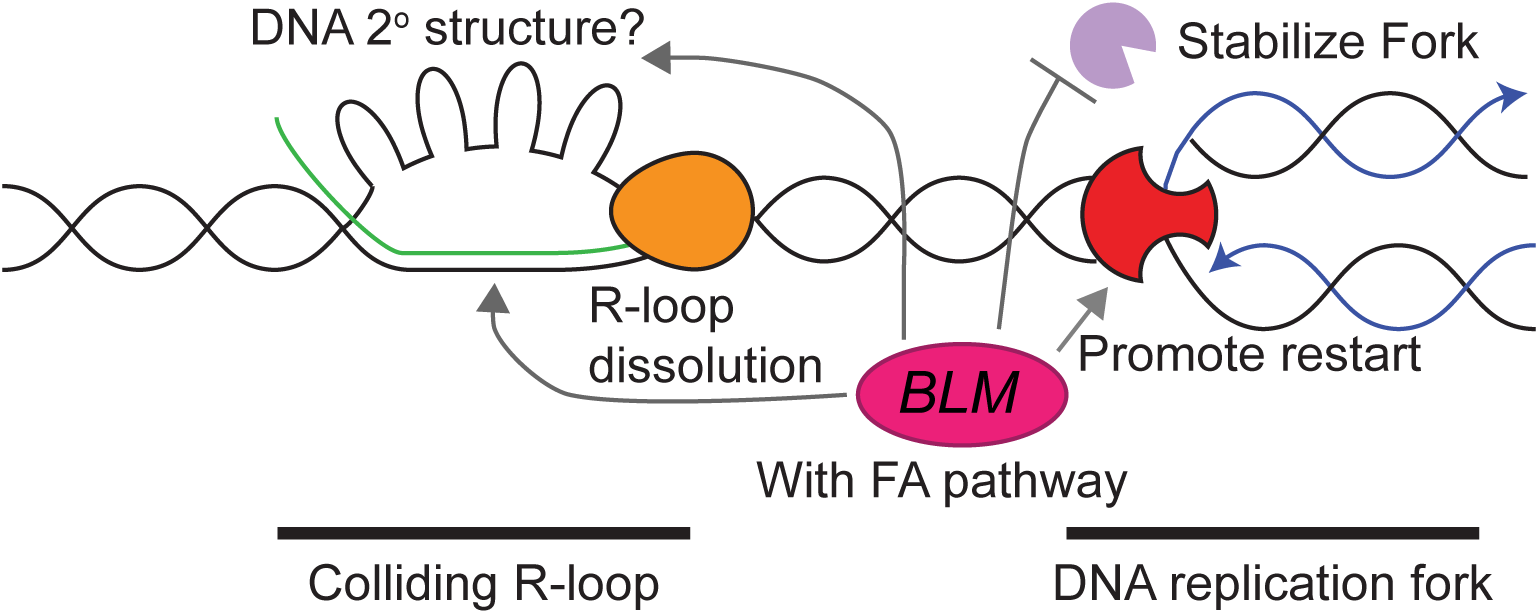
Model of Sgs1/BLM impact on R-loops. Shown are a replication fork (red) heading toward a stalled RNA polymerase (orange) and associated R-loop. Possible roles for BLM/Sgs1 at these sites are highlighted with arrows and discussed in the main text.

## DISCUSSION

### Mutation accumulation and whole genome sequencing

Here we use mutation accumulation and next generation sequencing to define the chromosome instability signatures of *SGS1* and *MUS81*-deficient cells. Loss of either gene increase large instability events much more than point mutations but the signatures are clearly distinguishable. Loss of Sgs1 leads primarily to copy number changes flanked by repetitive sequences, while Mus81 loss favors aneuploidy. These observations are consistent with the functions of each factor. Sgs1 rejects homeologous recombination events that may lead to CNVs between repeat elements (Myung et al., 2001). Mus81 cleaves concatenated chromosomes, which in the *mus81*Δ/Δ cells may remain attached during mitosis, lag and be segregated together to create an aneuploid daughter cell (Naim et al., 2013). Mutation accumulation approaches in model organisms have been powerful tools to understand mutation rates, and there are a growing number of studies that exploit the associated mutation signatures for mechanistic insights to mutagenesis in DNA replication and repair mutants and for gene-drug interactions (Meier et al., 2014; O’Connell et al., 2015; Segovia et al., 2017; Serero et al., 2014; Stirling et al., 2014). We believe these unbiased, genome-wide approaches will continue to be valuable to understand mutagenesis and gene-drug interactions and remain most accessible in model organisms.

### Mechanism of Sgs1/BLM action in preventing R-loop-driven genome instability

Our data show that loss of Sgs1 or BLM leads to R-loop accumulation across species and that at least some of the associated genome instability is contingent upon transcription-replication conflicts and/or R-loops. We find that in yeast the distribution of R-loops is increased at sites of Rrm3 occupancy and often linked to Sgs1 action, such as the rDNA, TY elements and telomeres, and also at a set of long genes. It is notable that reproducible differences in the DRIP-chip profile of *sgs1*Δ, occurred only at only some loci, and that the profiles were grossly similar to WT over the genome. Recently, it was observed that, despite binding them broadly, only a subset of R-loops are degraded by Rnh1 in yeast (Zimmer and Koshland, 2016). This highlights that there may be a poorly understood regulatory distinction between normal and abnormal R-loop formation in cells and we believe this distinction may be relevant to locus-specific differences seen in DRIP-chip profiles in mutant strains such as *sgs1*Δ. In the case of the rDNA, Sgs1 is known to have a role in facilitating both replication and transcription (Lee et al., 1999; Versini et al., 2003) and accordingly, we observed that enhanced rDNA instability seen in *sgs1*Δ cells is completely suppressed by RNaseH1 overexpression. In addition, we found that R-loops accumulated at long genes in *sgs1*Δ, and long genes have previously been identified as enriched for Rrm3 (Santos-Pereira et al., 2013), a helicase which facilitates replication through difficult to replicate regions. Indeed, Rrm3 occupancy correlated with increased DRIP signal in *sgs1*Δ. In addition, Top2, an interacting partner of Sgs1 which cooperates with Sgs1 and Top3 at rDNA (Mundbjerg et al., 2015), has also been linked to transcription at long protein coding genes in yeast (Joshi et al., 2012). Thus, regions that are sensitive to topological stress may be more likely to form R-loops in *sgs1*Δ cells.

Overall, we favor a model in which Sgs1/BLM works in proximity to R-loops at sites of replication-transcription conflict, consistent with studies of collaboration between Sgs1 and RNaseH2 (Chon et al., 2013; Kim and Jinks-Robertson, 2011). Within this direct model (**Fig. 7**) there remain several nonmutually exclusive possibilities: the simplest model is that Sgs1/BLM could unwind R-loops directly as supported by *in vitro* experiments ((Popuri et al., 2008) and **Fig. 6A**). Sgs1/BLM could also unfold G-quadruplexes, or other DNA secondary structures, associated with the non-template strand opposite to a DNA:RNA hybrid in a so-called G-loop (Duquette et al., 2004). The role of Sgs1/BLM in fork stabilization could also allow time for other factors to resolve R-loop blockages. Finally, Sgs1/BLM could direct the activity of another helicase as, at least in human cells, it is known to bind FANCM and FANCJ (Suhasini and Brosh, 2012). Indeed, the collaboration between BLM and FA pathway is very likely to be important for mitigating the effects of R-loops in human cells (Garcia-Rubio et al., 2015; Schwab et al., 2015). For example, BLM physically and functionally interacts with FANCM (Ling et al., 2016), potentially coordinating its activity with the activity of the FA pathway, and FANCM can use its strand migration activity to remove R-loops (Schwab et al., 2015). Indeed, FANCD2, which our data places in the same pathway as BLM for R-loop suppression, also regulates BLM stability and assembly at stalled replication forks, and reciprocally, FANCD2 activation requires BLM (Chaudhury et al., 2013; Panneerselvam et al., 2016). Thus there are multiple levels of regulation to be explored across systems in future experiments. Indeed, the scenario is likely to be more complex in human cells as BLM paralogs WRN and RECQL5 have both been linked to DNA:RNA hybrid metabolism in the test tube or in cells (Chakraborty and Grosse, 2010; Saponaro et al., 2014).

### Defective DNA repair proteins shifting the R-loop landscape

Our data add Sgs1/BLM to a growing list of DNA repair proteins that seem to work toward the error free resolution of R-loops and the prevention of deleterious transcription-replication conflicts. NER factors XPF and XPG appear to actively cleave DNA at or around R-loops to initiate a NER mechanism for R-loop clearance(Sollier et al., 2014). The Fanconi Anemia pathway, including BRCA1 and BRCA2, also suppress R-loop-mediated genome instability and likely do so through their replication fork protection functions (Bhatia et al., 2014; Chang and Stirling, 2017; Garcia-Rubio et al., 2015; Hatchi et al., 2015; Schwab et al., 2015). Indeed, there is now considerable evidence that DNA damage alone is sufficient to induce R-loops in mammalian cells (Britton et al., 2014; Tresini et al., 2015). The idea that normal robust DNA replication itself prevents R-loop accumulation is also gaining support, for example with recent studies showing effects of the MCM helicase or POLD3 in preventing DNA:RNA hybrid accumulation (Tumini et al., 2016; Vijayraghavan et al., 2016). The abundance of fork-protection factors emerging as R-loop regulators supports a generalized concept that functional DNA replication machinery is an important way to mitigate deleterious transcription-replication collisions. We speculate that there are common mechanisms at play in cancers experiencing abnormal replication stress and that transcription-mediated genome instability will play a role in tumor mutation accumulation.

## Materials and Methods

### Yeast growth and media

Yeast were cultured according to standard conditions in the indicated media at 30°C unless otherwise indicated. Growth curves were conducted in YPD media in 96-well plates using a TECAN M200. The area under the curve was used to compute expected and observed fitness values (Stirling et al., 2011; Stirling et al., 2012). A list of yeast strains and plasmids used in this study can be found in **Table S3**.

### Mutation accumulation and whole genome sequencing

Mutation accumulation experiments were conducted as described (Segovia et al., 2017). Single colonies from passage 40 were grown overnight in YPD to prepare genomic DNA by two rounds of phenol-chloroform extraction (Stirling et al., 2014). Whole genomes were sequenced using the Illumina HiSeq2500 platform and sequence files were deposited at the NCBI sequence read archive (http://www.ncbi.nlm.nih.gov/sra) (accession #: SRP094860 for *mus81*Δ*/*Δ and *sgs1*Δ*/*Δ genomes, and SRP091984 for WT genomes). Read quality control, alignment to UCSC saccer3, and variant calling was performed exactly as described (Segovia et al., 2017). CNVs were detected using an in house version of CNAseq (Jones et al., 2010) and Nexus copy number 7.5.2 (Biodiscovery Inc.). Variants were manually checked for read support using the Integrated Genomics Viewer (IGV).

### Recombination and genome instability assays

Recombination events in L, LYΔNS, LNA, pARSHLB-IN, pARSHLB-OUT, pARSCLB-IN, pARSBLB-IN, L-lacZ and GL-lacZ systems (kind gifts of A. Aguilera) were scored by counting Leucine positive (LEU+) colonies (Gomez-Gonzalez et al., 2009; Gomez-Gonzalez et al., 2011; Gonzalez-Aguilera et al., 2008; Herrera-Moyano et al., 2014). Recombination frequencies, and cell viability, were obtained from the average value of three tests performed with 3-9 independent transformants each as described(Stirling et al., 2012). In assays where yeast strains were transformed with a recombination and an over-expression vector, recombination and viability plates maintained both plasmids. To measure rDNA stability, yeast with *URA3* inserted into the rDNA locus were treated as for the recombination assays except that loss of *URA3* was measured by the frequency of 5′fluoroorotic acid resistant colonies (Wahba et al., 2011). Cell viability was measured by growing test strains on SC minus uracil (-URA) plates. To maintain plasmids all aspects of the rDNA instability assay were done on media lacking leucine. Finally, frequencies of chromosome III loss were quantified in *MATα* haploid knockout collection strains using the a-like faker assay as essentially as described (Ang et al., 2016). Graphing and statistical analyses were done in Graphpad (Prism).

### Yeast chromosome spreads and live cell imaging

Chromosome spreads were performed as previously described (Chan et al., 2014a). For each sample, at least 60 nuclei were visualized, and the nuclear fluorescent signal was quantified using ImageJ (Schneider et al., 2012). Each mutant was assayed in quadruplicate. For comparisons purposes, the S9.6 median fluorescence intensity of the wild type strain of each experiment was used for normalization. Mutants were compared to wild type by the unpaired t test.

For live cell imaging cells expressing Rad52-YFP were grown to logarithmic phase prior to any indicated treatments. Log-phase cells, treated or untreated, were bound to concanavalin-A coated slides and imaged on a Leica dmi8 inverted fluorescence microscope using the appropriate filter sets (Stirling et al., 2012).

### DRIP-chip analysis

DRIP-chip was generated and analyzed as described (Chan et al., 2014a). Complete datasets can be found at ArrayExpress: E-MTAB-5582. Data was normalized using the rMAT software (Droit et al., 2010). DRIP-chip profiles were generated in duplicate with averaged and quantile normalized data used for plotting and calculating average enrichment scores. For Ty elements, we averaged all probes whose start sites fell within the element’s start and end position, including the LTRs. The same was done for rDNA genes, telomeres, and Rrm3 peaks (coordinates derived from (Herrera-Moyano et al., 2014)). CHROMATRA plots were generated as described previously with genes aligned by their TSS and sorted by length (Hentrich et al., 2012). Average gene profiles were generating by averaging all probes that mapped to the genes of interest. Here, probes mapping to features of interest were split into 40 bins and probes matching to the 1500 bp of flanking sequences were split into 20 bins. Gene length average gene profiles were generated by splitting all genes into gene-length classes and averaging probes in 150 bp increments. To compare length of enriched genes in *sgs1*Δ, genes covered at least 50% by a DNA:RNA hybrid occupancy greater than 1.5 over the WT profile were tabulated, those genes appearing in both *sgs1*Δ DRIP replicates were analyzed for length compared to the genome average using a Mann-Whitney test (GraphPad).

### Cell culture and Transfection

HeLa cells were cultivated in Dulbecco’s modified Eagle’s medium (DMEM) (Stemcell technologies), while HCT116 was grown in McCoy’s 5A media, both supplemented with 10% fetal bovine serum (Life Technologies) in 5% CO_2_ at 37°C. For RNA interference, cells were transfected with siRNAs targeting BLM (*si-BLM*), FANCD2 (*si-FANCD2*), or Luciferase GL3 Duplex as a control (*si-Luc*) with Dharmafect1 transfection reagent (Dharmacon) according to manufacturer’s protocol and harvested 48 hours after the siRNA administration. siRNA sequences are: si-BLM, 5′-GCUAGGAGUCUGCGUGCCGA-3′; si-FANCD2, 5′-GGUCAGAGCUGUAUUAUUC-3′; and si-Luc, 5′-GUUACGCUGAGUACUUCGA-3′ (Blackford et al., 2015; Schwab et al., 2015). For experiments with overexpression of GFP or nuclear-targeting GFP-RNaseH1 (gift from R. Crouch), transfections were performed with Lipofectamine 3000 (Invitrogen) according to manufacturer’s instructions 24 hours after the siRNA transfections.

### Immunofluorescence

For S9.6 staining, cells were grown on coverslips overnight before siRNA transfection and plasmid overexpression. 48 hours post-siRNA transfection, cells were washed with PBS, fixed with ice-cold methanol for 10 minutes and permeabilized with ice-cold acetone for 1 minute. After PBS wash, cells were blocked in 3%BSA, 0.1% Tween 20 in 4X saline sodium citrate buffer (SSC) for 1 hour at room temperature. Cells were then incubated with primary antibody S9.6 (1:500) (Kerafast) overnight at 4°C. For HCT116, nucleolin was co-stained by co-incubating with anti-nucleolin (Abcam) at 1:1000. Cells were then washed 3 times in PBS and stained with mouse Alexa-Fluoro-568-conjugated secondary antibody (1:1000) (Life Technologies) for 1 hour at room temperature, washed 3 times in PBS, and stained with DAPI for 5 minutes. Cells were imaged on LeicaDMI8 microscope at 100X and ImageJ was used for processing and quantification of S9.6 intensity in images. Only GFP-positive cells were quantified, and micronuclei were counted in asynchronous cells from the same slides. For FANCD2 foci, the immunostaining were performed the same way except fixation with 4% paraformaldehyde for 15min and permeabilization with 0.2% Triton X-100 for 5 min on ice. Primary antibodies for FANCD2 (Novus) and rabbit Alexa-Fluoro-568-conjugated secondary antibody were all diluted 1:1000. Where indicated, cells were treated with DMSO or 2mM HU (Sigma) for 2 hours before fixing.

### Neutral Comet Assay

The neutral comet assay was performed using the CometAssay Reagent Kit for Single Cell Gel Electrohoresis Assay (Trevigen) in accordance with the manufacturer’s instructions. Electrophoresis was performed at 4°C and slides were stained with PI and imaged on LeicaDMI8 microscope at 20X. Comet tail moments were obtained using an ImageJ plugin as previously described (Mathew et al., 2014). At least 50 cells per sample were analyzed from each independent experiment.

### Western Blotting

Whole cell lysates were prepared with RIPA buffer containing protease inhibitor (Sigma) and phosphatase inhibitor (Roche Applied Science) cocktail tablets and the protein concentration were determined by Bio-Rad Protein assay (Bio-Rad). Equivalent amounts of protein were resolved by SDS-PAGE and transferred to polyvinylidene fluoride (PVDF) microporous membrane (Millipore), blocked with 5% skim milk in TBS containing 0.1% Tween 20 (TBST), and membranes were probed with the following antibodies: BLM (abcam), GAPDH (Thermo Scientific), FANCD2 (Novus), α-tubulin (Life Technologies). Secondary antibodies were conjugated to Horseradish Peroxidase and peroxidase activity was visualized using Chemiluminescent HRP substrate (Thermo Scientific).

### Proximity Ligation Assay

Cells were grown on coverslips, washed with PBS and fixed with 4% paraformaldehyde for 15min. After permeabilization with 0.2% Triton X-100 for 5 min, cells were blocked in 3%BSA, 0.1% Tween 20 in 4XSSC for 1 hour at room temperature. Cells were then incubated with primary antibody overnight at 4°C (1:500 rabbit BLM antibody (Sigma) as negative control; 1:200 mouse S9.6 antibody as negative control; 1:1000 rabbit BLM with 1:200 mouse S9.6, or 1:1000 rabbit 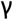H2AX (abcam) with 1:200 mouse S9.6 as positive control). After washing with 1XPBS twice, cells were incubated with pre-mixed PLA probe antimouse minus and PLA probe anti-rabbit plus (Sigma) for 1 hour at 37°C. Binding of PLA probes, ligation and amplification was performed with the reagents from the Duolink In Situ Kit (Sigma) according to manufacturer’s instructions. Slides were mounted in Duolink In Situ Mounting Medium with DAPI and imaged on LeicaDMI8 microscope at 100X.

### In vitro helicase assay

BLM was tagged at the N-terminus with GST and at the C-terminus with His_10_ and purified as described for PALB2 (Buisson et al., 2010). R-LOOP and D-LOOP substrates were generated by annealing purified oligonucleotides:

DNA strand 1: 5′GGGTGAACCTGCAGGTGGGCGGCTGCTCATCGTAGGTTAGTTGGTAGAATTCGGCAGCGTC-3′ DNA strand 2: 5′ -GACGCTGCCGAATTCTACCAGTGCCTTGCTAGGACATCTTTGCCCACCTGCAGGTTCACCC-3′ With either RNA: 5′-AAAGAUGUCCUAGCAAGGCAC-3′ or DNA: 5′-AAAGATGTCCTAGCAAGGCAC-3′. Unwinding assays were performed in MOPS buffer (25 mM MOPS (morpholine-propanesulfonic acid) pH 7.0, 60 mM KCl, 0.2% Tween-20, 2 mM DTT, 2 mM ATP, 2 mM MgCl2). BLM and labelled R-LOOP or D-LOOP (100 nM) substrates were incubated in MOPS buffer for 20 minutes at 37<u>°</u>C, followed by deproteinization in one-fifth volume of stop buffer (20 mM Tris-Cl pH 7.5 and 2 mg/mL proteinase K) for 20 minutes at 37<u>°</u>C. Reactions were loaded on an 8 % acrylamide gel, run at 150V for 120 minutes, dried onto filter paper and autoradiographed.

## ACKNOWLEDGEMENTS

We acknowledge Andres Aguilera, Allan Morgan, Frederic Chedin, Karlene Cimprich, Robert Crouch, Elizabeth Conibear, and Doug Koshland for providing reagents and methods. We thank Philip Hieter in whose laboratory some of the early work was conceived. This research is funded by the Canadian Cancer Society (grant 703263) and the Canadian Institutes of Health Research (CIHR) (MOP-136982) to P.C.S, CIHR project grant 363317 to J.Y.M and Natural Sciences and Engineering Research Council of Canada grant RGPIN 402095–11 to M.S.K. P.C.S. is a CIHR New Investigator and Michael Smith Foundation for Health Research Scholar. J.Y.M. is a Fond de Recherche du Québec Santé Chair in genome stability.

